# Dental erosion caused by a gastrointestinal disorder in a child from the Late Holocene of Northeastern Brazil

**DOI:** 10.1101/265702

**Authors:** Rodrigo E. Oliveira, Ana Solari, Sergio Francisco S. M. Silva, Gabriela Martin

**Author notes:** Corresponding author: Rodrigo E. Oliveira.

## Abstract

A skeleton of an approximately 3-years-old sub-adult, in an excellent state of conservation, was found at the Pedra do Cachorro rock shelter - Buíque, Pernambuco – Brazil, an archaeological site used as funerary place between 3875 and 575 cal years B.P. The skeleton has no signs of pathological bone changes, but its maxillary teeth show strong evidence of enamel and dentin wear caused by acid erosion, suggesting vomiting or gastroesophageal reflux episodes. The aim of this study was to describe the lesions and discuss the aetiology of these dental defects with the emphasis on the cause of the death of this individual.

## INTRODUCTION

In archaeological sites, teeth are usually found well preserved because they are made of the hardest tissues in the human body: enamel and dentin. They are generally used as a reliable source of information about physiological and pathological processes undergone by the individual (Ortner, 2003; Scheid and Weiss, 2012a).

In addition to dental diseases such as caries, periapical abscesses, dental calculus and periodontal resorption, dental wear is frequently used to determine diet, gastrointestinal diseases or parafunctional activities (d’Incau et al., 2012; Deter, 2009; Smith, 1984).

Dental wear can be determined by measuring mineral loss of organic tissues: enamel, dentin or both (Brace and Molnar, 1967; Molnar, 2008; Oliveira and Neves, 2015; Van’t Spijker et al., 2009). It is divided into four categories according to its causes: mechanical processes, such as abrasion, attrition and abfraction, or chemical processes, such as erosion. Abrasion is the most common cause of dental wear observed in archaeological human remains, whereas abfraction is less common (Grippo et al., 2012; Scheid and Weiss, 2012b). In fact, the loss of enamel and dentin is the result of two or more mechanical and chemical processes simultaneously (Lanigan and Bartlett, 2013; Lucas and Omar, 2012; Oliveira, 2014).

Some non-physiological activities can also change dental structure. Among others, parafunctional habits may include clenching and grinding (bruxism), which are very common in modern societies and may or may not be directly associated with psychosocial problems (Carlsson et al., 2003; Manfredini and Lobbezoo, 2009; Pavone, 1985). However, the most common parafunctional activity observed in prehistoric or traditional societies is the use of teeth as tools due to their resistance and the contraction strength of masticatory muscles, since teeth can grab and tear efficiently several material (Larsen et al., 1998; Waters-Rist et al., 2010).

The critical pH for hydroxyapatite solubilization (mineral structure of enamel) is 5.5, whereas the critical pH for fluorapatite (mineral formed from the reaction between hydroxyapatite and fluoride) is 4.5 (Ekstrand and Oliveby, 1999). Dental erosion is a loss of tooth structure in which an acidic agent is brought into contact with the surface of teeth, creating a microenvironment where pH remains below 4.0 (Dong et al., 1999; Hillson, 2008; Järvinen et al., 1991; Scheid and Weiss, 2012b). It is important to highlight that caries formed when biofilm produces lactic acid are not classified as dental erosion because in this case pH ranges from 5.5 to 4.0, whereas for dental erosion pH should be below 4.0(Larsen, 2008; Morimoto et al., 2014).

Some citrus juices and carbonated beverages which have a pH lower than 3.0 are responsible, nowadays, for most extrinsic erosion. In other words, excessive consumption of acid drinks can cause those changes on dental surfaces (Eccles and Jenkins, 1974; Honório et al., 2008; Järvinen et al., 1991; Lussi et al., 2011).

Some gastrointestinal diseases can also cause tooth erosion. Stomach acids can back up into the buccal cavity due to reflux and vomiting (Bartlett et al., 2013; Gudmundsson et al., 1995). Since gastric juice is extremely acid, with pH around 1.0, dental erosion can be used in the diagnosis of bulimia and other eating disorders (Gudmundsson et al., 1995; Lazarchik and Filler, 2000; Moazzez et al., 2004).

Dental erosion is often discussed in the medical and dental literature but is rarely described in the archaeological literature. This fact suggests that physical illnesses are responsible for the minority of cases of dental erosion, whereas mental disorders and the frequent ingestion of acidic food and beverages are the main etiological agents for dental changes in modern society (Dori and Moggi-Cecchi, 2014; Lanigan and Bartlett, 2013; Robb et al., 1991).

## CASE STUDY

The specimen analyzed in this study, which was named as Burial 2, was unearthed in 2015 from the Pedra do Cachorro archaeological site, located in the Parque Nacional do Catimbau, Pernambuco, Brazil (Figure 01). This site is a rock shelter that differs from the other archaeological sites of the region because there is no rock art in it. Between 2015 and 2016, four field campaigns were conducted at the site, resulting in the excavation of 68m^2^ Two burials were found during the first campaign. Burial 1 was a secondary burial of an adult male and Burial 2 was a primary burial of a sub-adult. In 2016, the secondary burial of an adult male was exhumed during the third campaign of excavation at the site (Solari et al., 2015, 2016).

**Figure 01.**
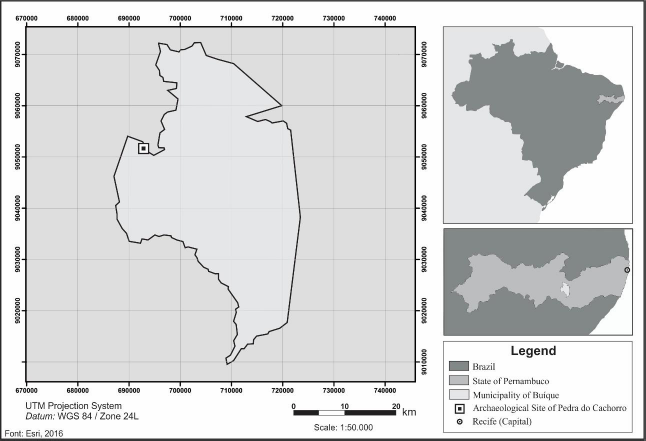
Location of the archaeological site of Pedra do Cachorro, Buíque - Pernambuco, Brazil.

Judging by the results of AMS dating of the rib bones of the three human skeletons, the Pedra do Cachorro site seems to have been used sporadically as a funeral area for 2800 years (Table 01).

**Table 01.**
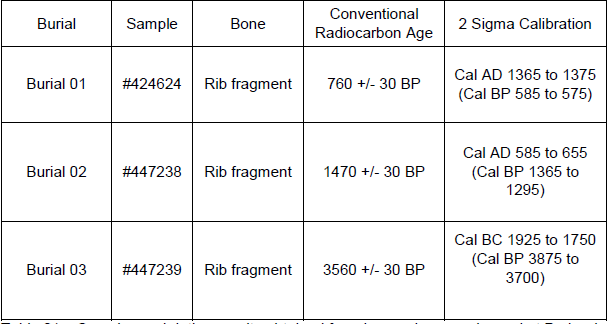
Samples and dating results obtained from human bones exhumed at Pedra do Cachorro site. Dating was carried out by Beta Analytic Laboratory.

Burial 2 was found in an oval pit (35cm of width, 92cm of length, and 20cm of depth) surrounded by sandstone blocks. The red-brown sediment of this pit slightly changed the color of this individual’s bones, although there is no evidence of intentional use of ochre or any other pigment on the body. A completely articulated skeleton was found, with no funerary offerings. Despite the large quantity of charcoal fragments associated with this skeleton, there was no macroscopic evidence of thermal changes in the child’s bones. This sub-adult had knees bent close to the chest, left arm extended along the body, right arm flexed next to the chest and hand on the face. The child was in prone position, oriented SW-NE (head-pelvis axis) (Figure 02) (Solari et al., 2016).

**Figure 02.**
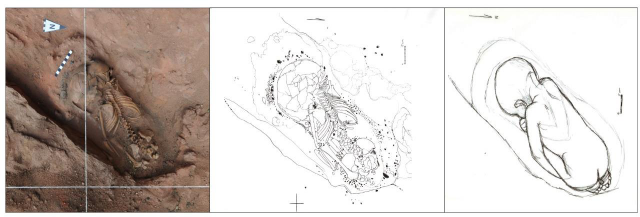
Burial 2: A. Photo of the exhumation; B. Burial sketch. C. Graphic reconstruction of the original burial position.

The well-preserved bones and very few *postmortem* fractures (observed only in the thin cranium and in the rib bones) allowed the interpretation of Burial 2 as a female who died when she was approximately 3 years old (± 1 year) (Figure 03).

**Figure 03.**
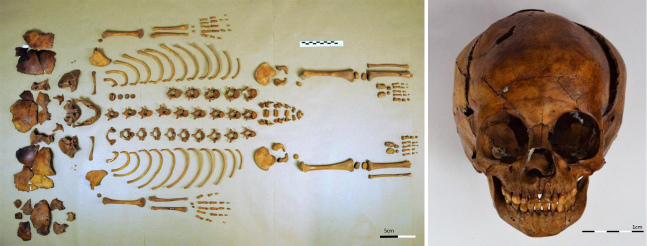
Skeleton and cranium from Burial 2 showing its excellent preservation.

The tooth development of this sub-adult was used for the age-at-death estimation. Sex determination was carried out considering the angle of the greater sciatic notch (Cunninghan et al., 2016; Schutkowski, 1993; Ubelaker, 1989). The well-preserved bones allowed us to measure the length of main upper and lower limbs and clavicle bones. Whereas the long bones suggest an age at death of around 1.5 years, the clavicle bones and the dental eruption of this sub-adult are compatible with a 3-year-old child (±1 year) (Table 02).

**Table 02.** Age estimation based on the length of bones of Burial 2.

| **Bones** | **Maximum Length (mm)** | **Estimated Age** | **References** |
| --- | --- | --- | --- |
| Femur L | 147,2 | 1,5 year | Maresh, 1970 |
| Femur R | 146,7 | 1,5 year | Maresh, 1970 |
| Tibia L | 121,9 | 1,5 year | Gindhart, 1973;<br>Maresh, 1970 |
| Tibia R | 120,7 | 1,5 year | Gindhart, 1973<br>Maresh, 1970 |
| Fibula L | 118,9 | 1,5 year | Maresh, 1970 |
| Fibula R | 120,4 | 1,5 year | Maresh, 1970 |
| Humerus L | 111,7 | 1,5 year | Maresh, 1970 |
| Humerus R | 110,9 | 1,5 year | Maresh, 1970 |
| Ulna L | 100,9 | 1,5 year | Maresh, 1970 |
| Ulna R | 99,5 | 1,5 year | Maresh, 1970 |
| Radio L | 92,2 | 1,5 year | Gindhart, 1973<br>Maresh, 1970 |
| Radio R | 91,3 | 1,5 year | Gindhart, 1973<br>Maresh, 1970 |
| Clavicle L | 66,0 | 2 to 3 year | Black and Scheuer,<br>1996 |
| Clavicle R | 64,9 | 2 to 3 year | Black and Scheuer,<br>1996 |

## ANALYSIS OF DENTAL CHANGES

Macroscopic observation was performed under artificial light to analyze the bones of the masticatory system and the teeth of Burial 2.

Regarding oral health, all deciduous teeth were present and correctly positioned. They also did not present dental calculus and bone resorption caused by periodontal disease. Both lower central incisors presented superficial caries in the cement-enamel junction (CEJ) on their labial surface. No dental abscesses, dental (enamel or dentin) hypoplasias or other dental pathological alterations were observed in both arches.

The lower teeth present occlusal wear resulting from attrition and/or abrasio;n compatible with degree 2 on Molnar’s scale for occlusal wear. The same degree of occlusal wear was observed in the maxillary teeth (Molnar, 1971; Smith, 1984) (Figure 04).

**Figure 04.**
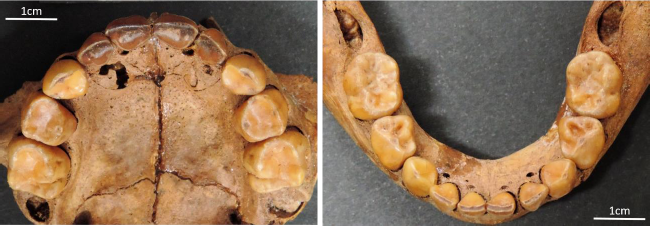
Maxillary and jaw from Burial 2. It is possible to observe physiological dental wear in all teeth and erosive changes in maxillary teeth.

Lingual surfaces of the deciduous upper molars (dm^1^) are very polished, and dentin was observed under a thin layer of enamel on the distolingual portion of dm^1^. The lingual surfaces of the maxillary canines are also very polished and the lingual surfaces of the four deciduous upper incisors show similar changes to those observed in the molars and deciduous canines, but in a more advanced stage. Lateral and central incisors show a quite strong dental wear bilaterally, also with substantial loss on the labial surface, exposing the dentin on the lingual surface of other four elements, and are classified as *IIIb* on the Eccles modified index for dental erosion (Eccles, 1979; Eccles and Jenkins, 1974). Despite the great loss of mineralized tissue, a thin enamel outline was observed on the lingual surface of the incisors (Figure 05).

**Figure 05.**
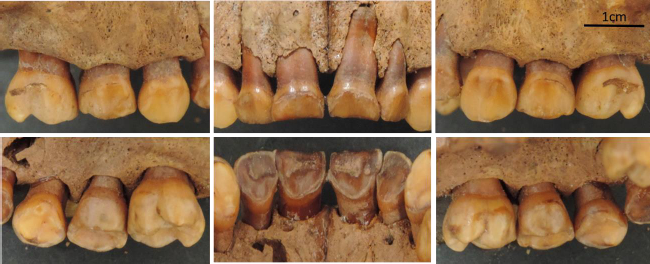
Detail of superior teeth (buccal/labial and lingual views), showing acid erosion on all incisors.

## DISCUSSION

Because of the low humidity of the region and the alkaline pH of the soil in which the child was buried, the skeleton from Burial 2 was very well-preserved (Baxter, 2004; Gordon and Buikstra, 1981; Manifold, 2015; Surabian, 2012).

This sub-adult had few carious lesions and no periodontal diseases, which is expected for a 3-year-old child. Her deciduous teeth could have been exposed for a short period to a cariogenic diet, such as breastfeeding, until the third year of life, as it was common in other prehistoric societies (Da-Gloria et al., 2017; Iida et al., 2007; Kaplan, 1996; Spielmann, 1989). This could explain her only two decays. The perfect position of lower and upper teeth indicates a satisfactory masticatory function, and the occlusal wear of teeth in both dental arches is associated with children who are mixed fed/weaned or who started exclusively masticatory function earlier in life (Martinez-Maza et al., 2016; Moynihan, 2005; Warren et al., 2002).

Parafunctional habits and intrinsic or extrinsic acid erosions are listed as possible factors that caused Burial 2 dental wear. Bruxism is one of the most common parafunctional habits that causes mineral loss. However, this parafunction could not have generated the abrasion angles observed in this case due to the limited movements of temporomandibular joint (TMJ) and, moreover, the lower teeth do not present any dental change compatible with and common in bruxism (Brace and Molnar, 1967; Molnar, 1971). Another possible parafunctional activity would be the daily use of teeth as tools for creating artifacts from vegetable fibers, leather, bones, and others. However, neither the lower teeth marks nor the maxillary incisor wear surface is consistent with this usage, not to mention that the child of Burial 2 was less likely to participate in these socio-cultural activities (Oliveira, 2014; Larsen et al., 1998; Molnar, 1971).

The maxillary incisors of Burial 2 show changes in buccal surfaces due to incisal surface loss. The buccal surfaces of canines, maxillary molars, and all mandibular teeth were not affected. Recurrent episodes of vomiting or chronic reflux caused a strong demineralization of the lingual surface of anterior maxillary teeth and moderate to slight demineralization of posterior maxillary teeth due to the posterior-anterior direction of the flow of gastric fluids into the mouth (Bartlett et al., 2013; Lazarchik and Filler, 2000). The buccal surface of maxillary teeth is partially protected by the oral mucosa, whereas mandibular teeth are protected by the cheek and tongue during vomiting, protecting these dental surfaces from gastric fluids, as observed in Burial 2 (Linnett and Seow, 2001). In fact, abrasion and attrition may have contributed to this scenario, but the evidence presented here shows acidic erosion and does not show other processes (Grippo et al., 2012; Lanigan and Bartlett, 2013; Lussi et al., 2011).

Finally, there are several diseases that can cause vomiting or gastroesophageal reflux and, consequently, dental erosion, but few or none can cause bone changes. Some diseases, such as gastrointestinal inflammatory diseases, anatomical abnormalities, malignant tumors, intracranial hypertension, central nervous system infection, metabolic diseases, and toxic food intake, among others, may have caused gastric disorders (Katz et al., 2013; Nebel et al., 1976; Rudolph et al., 2001; Vakil et al., 2006; van Herwaarden et al., 2000; Vandenplas et al., 2009). A gastrointestinal disorder or systemic disease could be associated with malnutrition in the child and could explain the small size of her long bones. These chronic disorders could be associated with the premature death of this child (Deaton, 2008; Kielmann and McCord, 1978; Maitland et al., 2006; Onis, 2010; Rice et al., 2000; van den Broeck, 1995).

## CONCLUSION

A well-preserved skeleton of a 3-year-old child exhumed during an archaeological excavation in northeastern Brazil, dated to 585-655 AD, was used as a case study to obtain evidence of gastrointestinal illness that causes severe dental erosion in maxillary incisors.

Despite the difficulties in diagnosing the specific gastrointestinal diseases that caused the dental erosion, it is important to accurately describe this case to show that acidic dental erosion also happened at that time. This study helps us to better understand the quality of life of pre-colonial inhabitants of semi-arid regions in northeastern Brazil.

Further chemical and biomolecular analysis should be carried out to find out the child’s cause of death.

## ACKNOWLEDGMENTS

The authors are grateful for the institutional support of the Departamento de Arqueologia da Universidade Federal de Pernambuco (DARQ-UFPE) and the financial support of the Instituto Nacional de Ciência e Tecnologia de Arqueologia, Paleontologia e Ambiente do Semiárido do Nordeste do Brasil (INCT-INAPAS). Ana Solari thanks the financial support of CAPES-PNPD.

